# Can school children support ecological research? Lessons from the *‘Oak bodyguard’* citizen science project

**DOI:** 10.1101/712638

**Authors:** Bastien Castagneyrol, Elena Valdés-Correcher, Audrey Bourdin, Luc Barbaro, Olivier Bouriaud, Manuela Branco, György Csóka, Mihai-Leonard Duduman, Anne-Maïmiti Dulaurent, Csaba B. Eötvös, Marco Ferrante, Ágnes Fürjes-Mikó, Andrea Galman, Martin M. Gossner, Deborah Harvey, Andy G. Howe, Michèle Kaennel-Dobbertin, Julia Koricheva, Gábor L. Löveï, Daniela Lupaștean, Slobodan Milanović, Anna Mrazova, Lars Opgennoorth, Juha-Matti Pitkänen, Marija Popović, Tomas V. Roslin, Michael Scherer-Lorenzen, Katerina Sam, Marketa Tahadlova, Rebecca Thomas, Ayco J. M. Tack

**Affiliations:** BIOGECO, INRA, Univ. Bordeaux, 33610 Cestas, France; Dynafor, INRA-INPT, University of Toulouse, Auzeville, France; CESCO, Museum National d’Histoire Naturelle, CNRS, Sorbonne-University, Paris, France; ”Ștefan cel Mare” University of Suceava, Forestry Faculty, Applied Ecology Laboratory, Universității Street 13, Suceava, Romania; Centro de Estudos Florestais, Instituto Superior de Agronomia, Universidade de Lisboa; NARIC Forest Research Institute, Department of Forest Protection, Hegyalja str. 18, 3232 Mátrafüred, Hungary; UniLaSalle, AGHYLE, UP.2018.C101, SFR Condorcet FR CNRS 3417, FR-60026 Beauvais, France; Department of Agroecology, Aarhus University, Flakkebjerg Research Centre, DK-4200 Slagelse, Denmark; Mitrani Department of Desert Ecology, Ben-Gurion University of the Negev, Midreshet Ben-Gurion, 8499000, Israel; Misión Biológica de Galicia (MBG-CSIC), Pontevedra, Galicia, Spain; Forest Entomology, Swiss Federal Research Institute WSL, Zürcherstrasse 111, CH-8903 Birmensdorf, Switzerland; School of Biological Sciences, Royal Holloway University of London, Egham, UK TW20 0EX; Department of Geosciences and Natural Resource Management, University of Copenhagen, Denmark, Rolighedsvej 23, 1958 Frederiksberg C, Denmark; University of Belgrade, Faculty of Forestry, Kneza Višeslava 1, 11000 Belgrade, Serbia; Mendel University, Faculty of Forestry and Wood Technology, Zemedelska 3, 61 300 Brno, Czech Republic; Biology Centre CAS, Entomology Institute, Ceske Budejovice, 37005, Czech Republic; University of South Bohemia, Faculty of Science, Ceske Budejovice, 37005, Czech Republic; Swiss Federal Research Institute WSL, Biodiversity and Conservation Biology, Ecological Genetics, Zürcherstrasse 111, 8903 Birmensdorf; Department of Ecology, Philipps-Universität Marburg, Karl-von-Frisch Strasse 8, 35043 Marburg; Department of Ecology, Spatial Foodweb Ecology Group, Department of Agricultural Sciences, PO Box 27 (Latokartanonkaari 5), FI-00014 University of Helsinki, Finland; Geobotany, Faculty of Biology, University of Freiburg, Schaenzlestr. 1, 79104 Freiburg, Germany; Department of Ecology, Environment and Plant Sciences, Stockholm University, SE-106 91 Stockholm, Sweden

## Abstract

Scientific knowledge in the field of ecology is increasingly enriched by data acquired by the general public participating in citizen science (CS) programs. Yet, doubts remain about the reliability of such data, in particular when acquired by school children. We built upon an ongoing CS program - *Oak bodyguards* - to assess the ability of European schoolchildren to accurately estimate the strength of biotic interactions in terrestrial ecosystems. We used standardized protocols to estimate predation rates on artificial caterpillars and insect herbivory on oak leaves and compared estimates made by school children, trained and untrained professional scientists (with no or limited expertise in predation or herbivory assessment). Compared to trained scientists, both schoolchildren and untrained professional scientists overestimated predation rates, but assessments made by the latter were more consistent. School children overestimated insect herbivory, as did untrained professional scientists. Thus, raw data acquired by school children participating in CS programs cannot be used and require several quality checks. However, such data are of no less value than data collected by untrained professional scientists and can be calibrated for bias.

## Introduction

Scientific knowledge is more accessible than ever before, particularly due to an increase in open-access publications and outreach activities of scientists worldwide. Still, many science topics in life and environmental sciences are highly controversial in society, even among individuals with substantial science literacy and education^1–3^. Citizen science (CS) programs rely on participation of the general public in scientific research in collaboration with or under the direction of professional scientists^4,5^. The rapid development of these programs, in addition to vastly increasing available data, offers an unprecedented opportunity to bridge gaps between science and society, by engaging the general public with the process of science, and increasing motivation and interest in scientific topics.

Citizen science programs in the field of ecology can benefit both science and society^6^. For professional scientists, involving the general public enables the collection of data on broader spatial and temporal scales than would otherwise be possible (*i.e.*, ‘crowdsourcing’). This practice has been recognized as a highly effective way to track various biological phenomena^7,8^. Typical CS studies in ecology address the effect of environmental factors on biodiversity^e.g. 9–11^ or climate change impact on plant or animal phenology^8,12,13^. In turn, volunteers engaged in CS programs can gain recognition for their skills and develop a deeper understanding of scientific concepts and the scientific process^14^. This may positively contribute to both science and environmental education^6^ and raise awareness of environmental issues. As a result, CS programs are now promoted by major funding agencies in Europe and North America^e.g., 4,,15^.

Engaging school children and their teachers can enhance the long-term educational and social goals of CS programs, for several reasons^16^. First, school pupils are guided by their instructors when learning about the scientific question raised by the CS program, as well as about the nature and social aspects of science^17,18^. Second, exposure to outdoor nature during childhood increases motivation and provides a long-lasting positive relationship with the environment while increasing people’s knowledge about nature^19,20^. Third, targeting schoolchildren for CS projects has the potential to engage a wider cross-section of society in science ^22^ than other CS projects involving self-selecting volunteers (they choose whether to be involved which can lead to the underrepresentation of many social groups, although strategies exist to increase engagement^21^).

Nonetheless, the enthusiastic views of win-win interactions through CS programs have been questioned by social scientists and ecologists^22^. The former point out that the educational and social impact may be overstated^14,23–26^, while the latter are concerned about the accuracy of data collected by the general public^27^, especially when school children are involved. The main reason for these concerns is that CS data are arguably of lower quality than those collected by professional scientists^16,24,27^. It has been proposed that data collected by school children involved in CS programs can contribute to environmental research, provided that research methods are kept simple and require skills that the children already have or are able to gain when mentored by adults^10,11,16^, and that the participant receives training, even remotely^28^. However, only a few studies have directly compared the quality of data acquired by professional scientists *vs.* the general public^10,11,29^. Quantitative evaluations of the impact of CS programs on science are thus needed to reciprocally engage citizens with science, and scientists with citizens.

Here, we report on the preliminary results of the ‘*Oak bodyguards*’ CS program which has so far involved school children and professional scientists in 14 European countries. The project aims to assess the effects of climate on two key biotic interactions occurring widely in natural and anthropogenic ecosystems, *i.e.*, the top-down and bottom-up forces controlling insect herbivory on leaves of the English oak (*Quercus robur*). We chose the English oak as a model species as it is one of the most common and emblematic forest trees in Europe, with a geographic range spanning more than 19 degrees of latitude. Furthermore, it is also widespread in both natural, rural, suburban and urban environments. In this project, school children and professional scientists exposed dummy plasticine caterpillars in oak trees to estimate predation rates^30–32^. We assessed the accuracy of CS data by comparing predation rate and insect herbivory estimates by three types of observers: professional scientists with previous experience in the project methodology (‘*trained professional scientists*’ henceforth), professional scientists with no previous experience in the project methodology (‘*untrained professional scientists*’), and schoolchildren. We first compared caterpillar predation rate estimates by schoolchildren or professional scientists (trained and untrained) with those of single scientist (Elena Valdés Correcher, EVC) used a control. Second, in a separate experiment, schoolchildren, trained and untrained professional scientists estimated leaf insect herbivory as the percentage of leaf area removed or damaged by insect herbivores^33^. We compared herbivory estimates by these three categories of observers. Our study asked whether schoolchildren were able to conduct an ecological experiment and acquire scientific data of a quality comparable to that acquired by professional scientists. We use the results to discuss risks and opportunities for the future of CS programs with schoolchildren.

## Results

### Predation rate

In total, 5,520 dummy caterpillars were installed on 153 oak trees by 34 partner schools in eight countries and 28 scientific partners in 14 countries throughout Europe **(Figure 1)**. Dummy caterpillars were exposed on trees for 16 days on average (range: 3 - 56). Among the 1,775 dummy caterpillars installed by school children, 640 were identified by EVC with attack marks by predators (*i.e.*, 36.06 %). Among the 3,745 dummy caterpillars installed by professional scientists, 1,268 were found to be attacked by predators (33.86 %).

**Figure 1.**
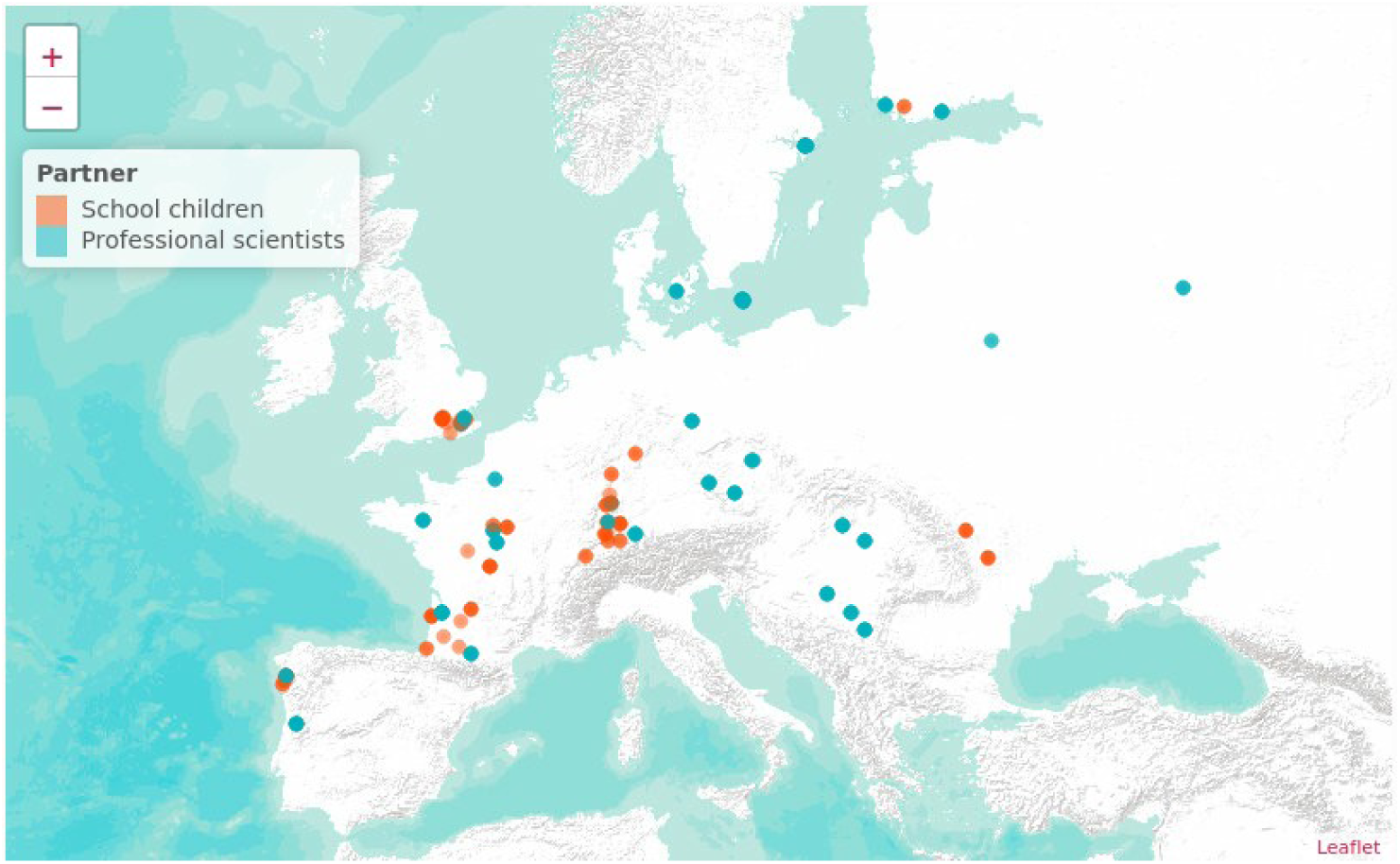
Location of oak trees included in the study. See the supplementary material for an interactive version.

Both school children (paired *t*-test: *df =* 38, *t* = −6.31, *P* < 0.001) and professional scientists (*df* = 158, *t* = - 3.80, *P* < 0.001) overestimated predation rate as compared to estimates made by a single trained observer **(Figure 2)**. Detailed examination of pairwise comparisons at the tree level confirmed that partner schools consistently overestimated predation rates as compared to a single observer, while the sign of deviation in scoring by scientific partners was more balanced between over- and under-estimation **(Figure 2)**.

**Figure 2.**
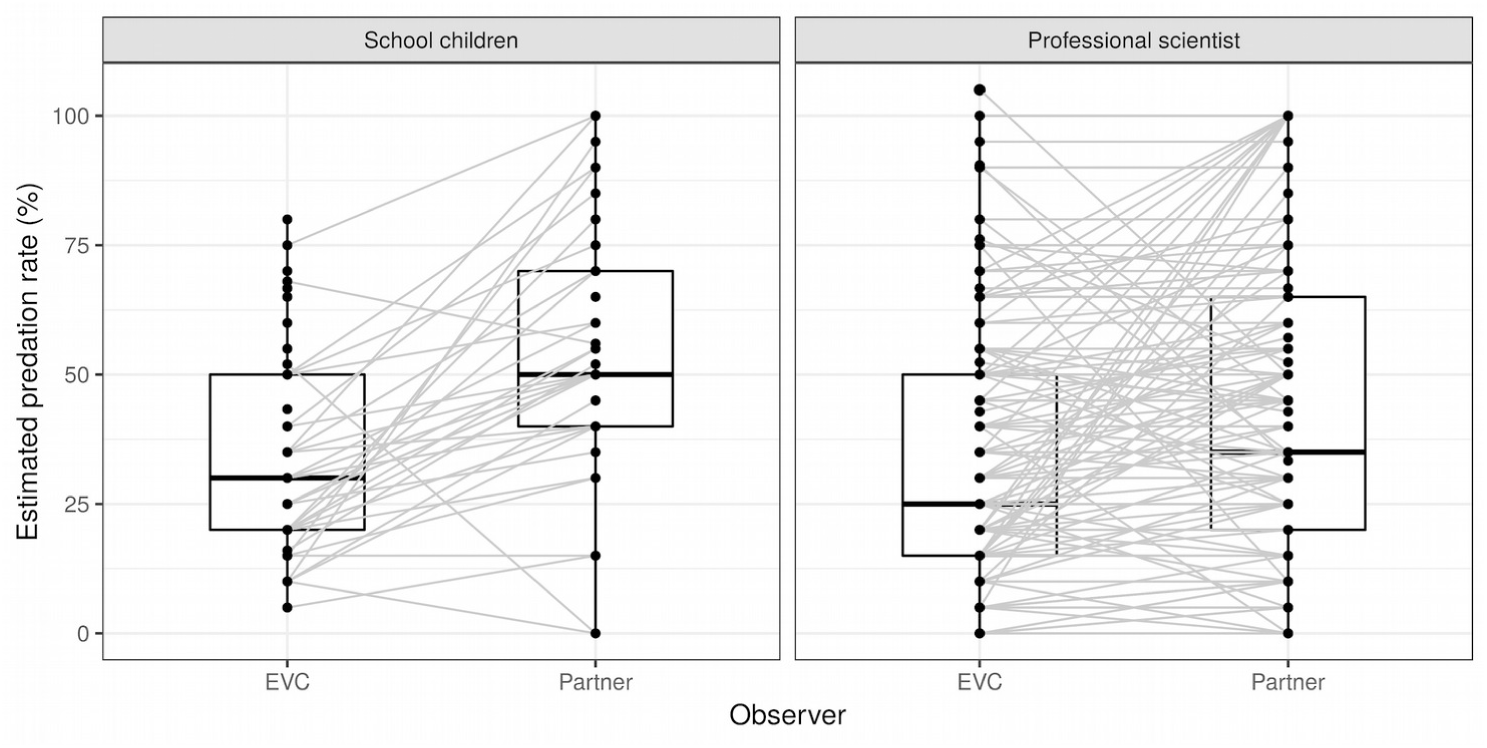
Comparison of predation rate estimated by school children or professional scientists *vs*. a single trained observer (Elena Valdés-Correcher). Points joined by grey lines represent the same tree.

Predation rate estimates by school children were more biased (intercept estimate ± SE: β_0_ = 37.29 ± 7.92) than those by professional scientists (β_0_ = 18.08 ± 3.16). Likewise, school children made less accurate predation assessments (slope estimate ± SE: β_1_ = 0.42 ± 0.23) than professional scientists did (β_1_ = 0.66 ± 0.06, **Figure 3**).

**Figure 3.**
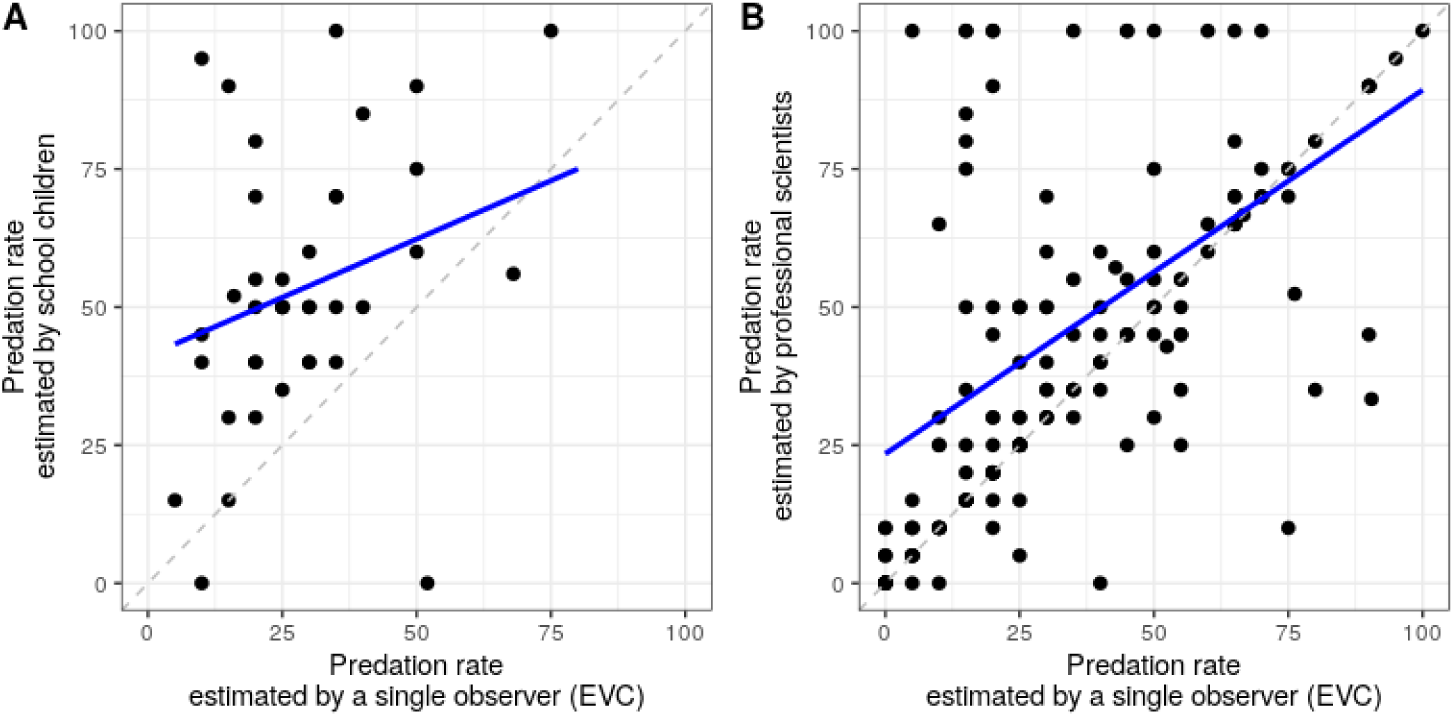
Precision and accuracy of school children and professional scientists (B) in assessing predation rate (% artificial larvae with predation marks). Dots represent predation rate aggregated at the level of oak trees for each survey separately. Dashed lines indicate a 1:1 relation. Bold blue lines represent linear regression slopes of predation rate by school children or professional scientists on assessments made by a single trained observer (EVC: Elena Valdés-Correcher). Regression equations: y = 0.42·x + 41.16, marginal (fixed effects) R_m_^2^ = 0.07, conditional (fixed *plus* random effects) R_c_^2^ = 0.53 **(A)** and y = 0.66·x + 23.42, R_m_^2^ = 0.31, R_m_^2^ = 0.79 **(B)**.

### Insect herbivory

Insect herbivory estimates by trained professional scientists were the lowest (mean ± SE = 9.00 ± 0.51 %, range 2.20 to 19.6 %) **(Figure S2)** whereas insect herbivory estimates by untrained professional scientists were the highest (14.65 ± 1.01 %, range from 3.80 to 62.00) **(Figure S2)**. Schoolchildren estimates of insect herbivory were intermediate (11.55 ± 0.64 %, range from 2.20 to 27.40 %) **(Figure S2)**. Thus, both untrained professional scientists and school children consistently overestimated insect herbivory compared to trained professional scientists (*F*_2,22_ = 27.31, *P* < 0.001) **(Figs 4 and S2)**. Interestingly, untrained professional scientists overestimated insect herbivory compared to both school children and trained professional scientists **(Figure 4)**.

**Figure 4.**
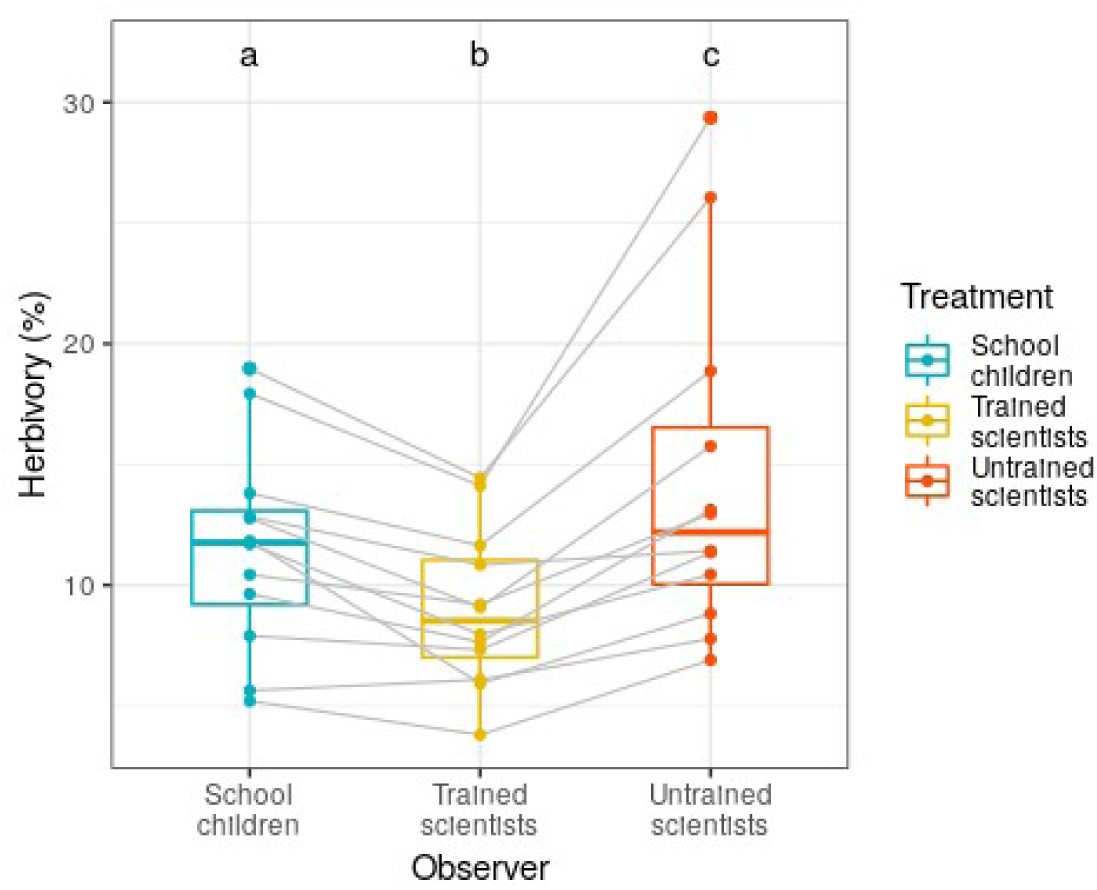
Pairwise comparisons between insect herbivory as estimated by school children, trained and untrained professional scientists. Boxplots represent mean insect herbivory aggregated across observers at the level of individual leaf sets. Thick grey lines connect dots corresponding to the same leaf sets. Different letters above boxplots indicate significant differences between treatments.

Herbivory estimated by school children was less biased (β_0_ = 0.84 ± 1.58) than that of untrained professional scientists (β_0_ = −3.5 ± 2.61). Likewise, school children made more accurate herbivory assessments (β_1_ = 1.19 ± 0.17) than untrained professional scientists did (β_1_ = 1.99 ± 0.27, **Figure 5**). Note that on average, unlike school children, untrained professional scientists overestimated herbivory across the observed range of herbivory. The negative intercept for untrained professional scientists here is driven by a slope estimate close to β_1_ = 2.

**Figure 5.**
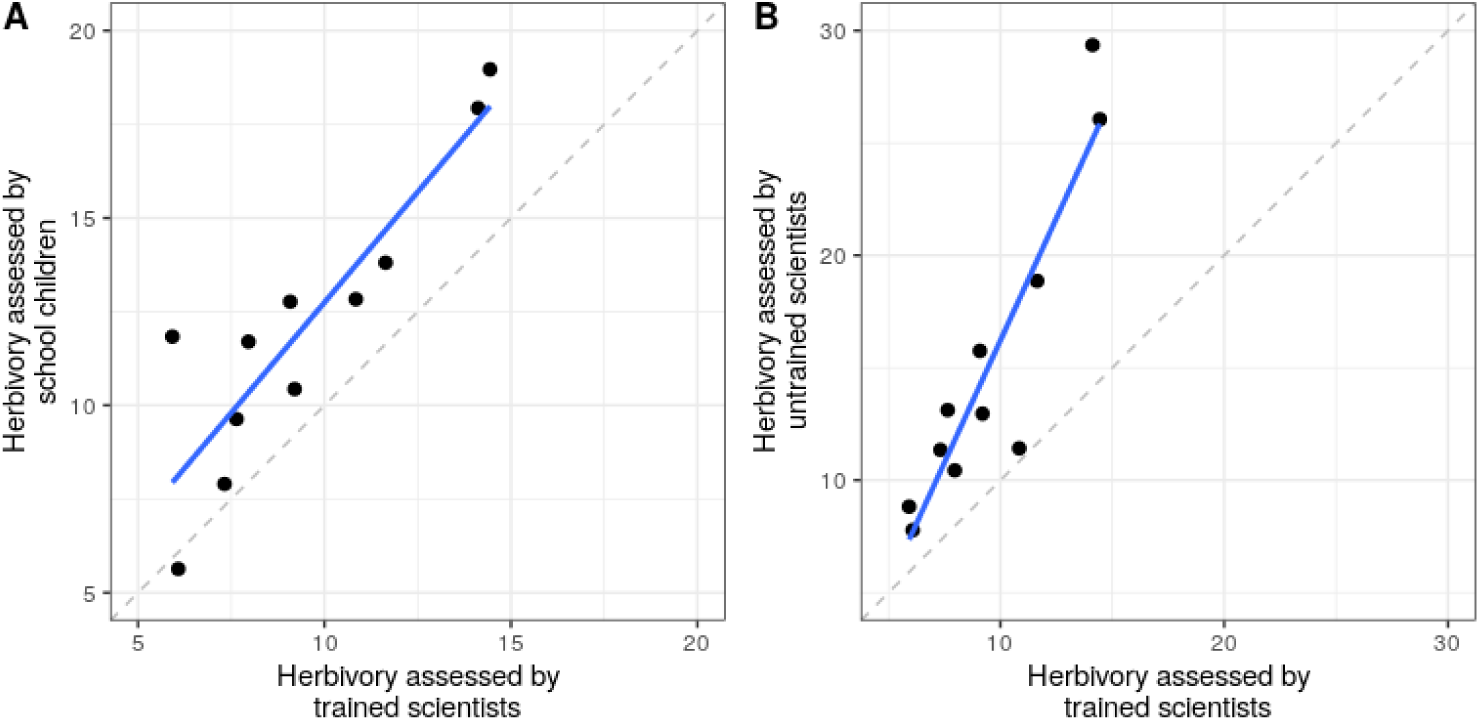
Precision and accuracy of school children and untrained professional scientists in assessing insect herbivory. Note the x- and y-axes are on the same scale in each panel. Dots represent mean insect herbivory aggregated across observers at the level of individual leaf sets. Dashed lines indicate a 1:1 relation. Bold blue lines represent linear regression slopes of herbivory assessed by school children or untrained professional scientists on assessments made by trained professional scientists. Regression equations: y = 1.19·x + 0.84, R^2^ = 0.84 **(A)** and y = 1.99·x - 3.5, R^2^ = 0.84 **(B).**

## Discussion

Our comparison of data collected by different audiences (schoolchildren, untrained and trained scientists) allowed us to examine the quality of ecological data collected by schoolchildren, to identify factors causing differences in the accuracy of data collected by different types of participants, and to suggest improvements for future CS programmes.

### Can school children collect data of relevant quality for ecological research?

The main strength of CS programs, from a research perspective, is the collection power achieved by volunteers (especially if the data are independently verified). However, our findings proved ambivalent with respect to whether the resulting data are of sufficient quality to yield scientifically robust results. On one hand, we clearly show that school children overestimated both predation rate and insect herbivory as compared to trained professional scientists. On the other hand, professional scientists with mixed expertise in these fields *also* tended to overestimate predation rate and clearly overestimated insect herbivory. Importantly for the interpretability of the data, overestimation was consistent across partner schools, as overestimation occurred in 35 observations (out of 39, *i.e.* 90%). Predation rates as assessed by professional scientists were, on average, slightly higher than predation rates (re-)estimated by a single trained observer. However, pairwise comparisons revealed that overestimation occurred in only 40% of observations (*n* = 65). Collectively, our results indicate that data provided by school children should be considered with caution, but the same holds true for data provided by untrained professional scientists.

### Why did (so) many school partners overestimate predation rate?

Overestimation principally arose from partners scoring scratch marks left by contact with buds or leaves as signs of predation **(Figure 6)**. Although the protocol mentioned this type of mark could be seen and should not be counted as predation, we did not include photos in the first version of the field bite guide. Other sources of overestimation of predation cannot be ignored. Although no teachers mentioned vandalism of experiments, researchers should be aware of this possibility, particularly when caterpillars are exposed on trees in urban environments. This may lead to ‘missing’ caterpillars falsely scored as attacked. In addition, school children were told by teachers that the aim of the study was to determine “who protects oaks” against herbivores. It is possible that school children (and their teachers too) felt they *had to* see predation marks, because this is what they perceived as the aim of the experiment. However, although confirmation bias is more likely to occur in school children and their teachers, it is important to stress that this type of cognitive bias is also common among trained professional scientists, who may have interpreted e.g. small cracks on caterpillar surface as predation marks^34,35^.

**Figure 6.**
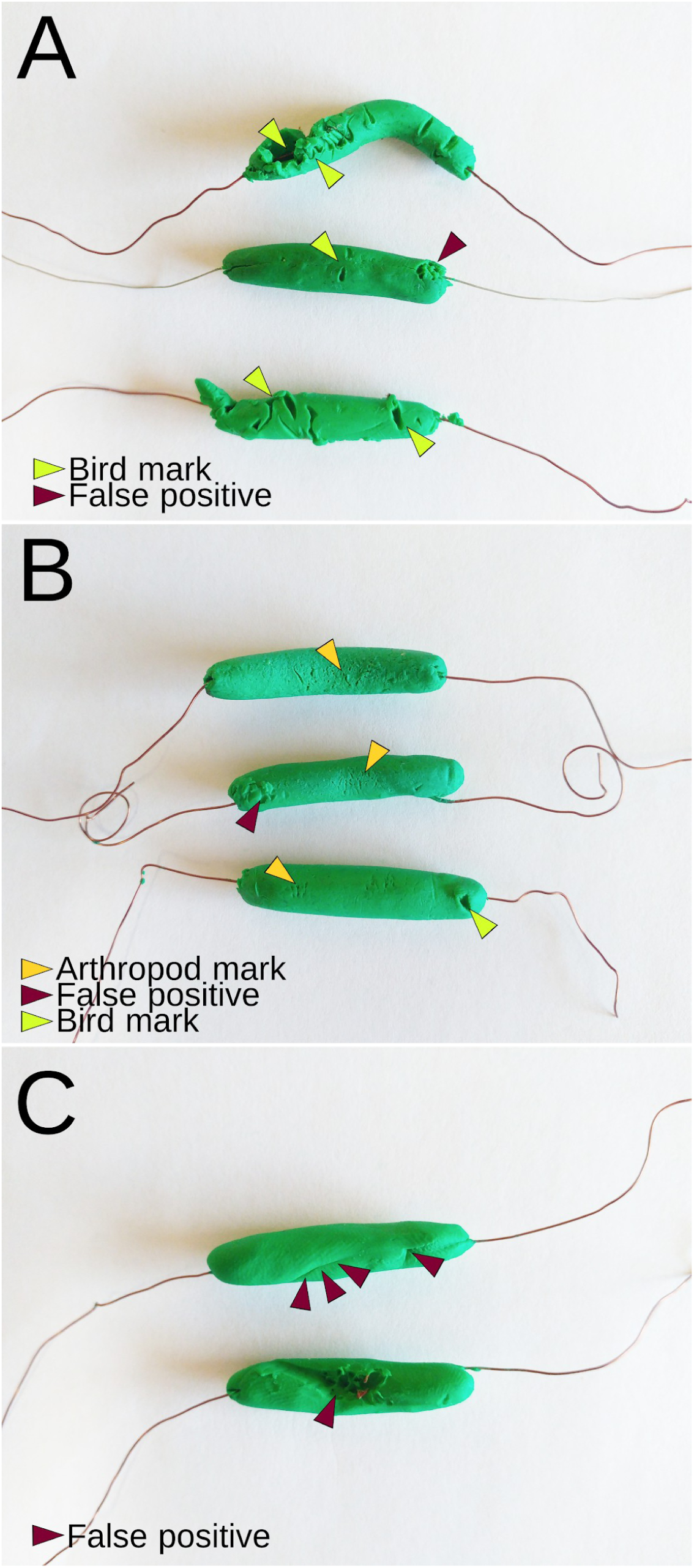
Examples of real and false positive observations of predations. In **A**, green arrows point to typical bird predation marks. The brown arrow points toward marks made by the wire when attaching the caterpillar on the branch and taking it off. In **B**, yellow and green arrows indicate marks made by arthropod mandibles and bird beaks, respectively. Brown arrows point to a beak mark. In **C**, brown arrows indicate typical marks erroneously counted as predation marks by school children. The scar-like mark on the top caterpillar was made when rolling the caterpillar onto the wire. Deep marks on the bottom caterpillar are imprints of branches and buds.

Although the protocol clearly specified how to standardize caterpillar size and shape, and emphasized the importance of standardization, we noticed that the dimensions of dummy caterpillars varied widely, both within and among partner schools. In other studies, the probability of detecting predation marks left by avian or arthropod predators was found to be influenced by the length and width of artificial caterpillars^32^. Unfortunately, we were unable to quantify the dimensions of caterpillars from the images we received. It is unlikely that variability in the dimension of artificial caterpillars has affected the comparison of predation rate as estimated by school children *vs*. trained professional scientists. However, the variation found should be regarded as a potential source of bias in large-scale multi-partners studies. As a potential mitigation procedure, researchers can provide pre-made caterpillars to project partners^31^. However, making caterpillars according to a standard protocol is also an important dimension of student training. At least, scientists should provide partners with a reference caterpillar made of hardened undeformable clay. 3D-printed models of caterpillars attacked by different predator types may also be included as examples. In any case, we advise that project partners be instructed to carefully pack caterpillars when sending these to lead scientists for calibration of predation assessment. We also recommend that data collected by school children are not directly used in the project, but undergo verification by trained professional scientists.

### School children overestimated insect herbivory; untrained professional scientists did too

Johnson et al.^33^ found that bias in herbivory assessment decreased with the number of years of experience in herbivory assessment. Interestingly, in the present study, variability in herbivory aggregated at the level of individual leaf sets was lower when it was estimated by trained professional scientists or school children than when herbivory was estimated by untrained professional scientists **(Figure S2)**. This result is consistent with the observation that although school children overestimated herbivory **(Figure 5)**, their estimates were more accurate than those made by untrained professional scientists **(Figure 5)**. It is possible that school children took the activity more seriously than untrained professional scientists did. An alternative explanation for this unexpected finding is that school children formed groups of 2-3 participants, while untrained professional scientists were alone when estimating herbivory. Within-group discussion may have reduced subjectivity and variability of estimates made by school children. Regardless of the cause, these results clearly indicate that school children are no less reliable than untrained professional scientists when it comes to estimating insect herbivory (on oak leaves).

### How can we make data collected by school children more reliable?

Citizen science programs can help to generate a large amount of data, but the quality has been questioned, especially when these ‘big data’ are not well structured by standard protocols^27,36^. Few studies have evaluated the quality of data collected by school children participating in citizen science programs^10,11,29^. It emerges from these studies that school children can actually provide data accurate enough to support ecological research, provided that the tasks they are requested to undertake are adapted to their skills and that they receive proper training^10,11,28^. Although we could not provide face-to-face training sessions for every school partner involved in the ‘*Oak bodyguards*’ project, the project methodology was simple and based on a detailed protocol. Nonetheless, this simplicity did not suffice to guarantee unbiased data, as illustrated by the fact that schoolchildren consistently overestimated predation rates. We therefore emphasize that citizen science programs relying on data collected by school children should include several checks of data quality and appropriate mitigation procedures. In particular, training sessions undertaken face-to-face or at least remotely must be planned before data collection^28^. Finally, whenever possible, the researcher analyzing the data should recover the raw material collected by children, or at the very least pictures allowing the re-assessment of measurements^12,29^. Importantly, these recommendations also hold true for large multi-partners research programs, as we also detected bias in data collected by professional scientists^34^. Whether variability in observations made by school children is random or can be modelled using appropriate covariates is an important question deserving further attention.

### Conclusion

Here we found that school children involved in CS programs can support ecological research, but only if their contributions are considered with caution. The acquisition of reliable data requires experimental procedures that are easy to implement, but even so, a measurement of interpretation bias seems essential. Several quality checks and curation procedures are therefore needed prior to using data collected by school children for ecological research. Unexpectedly, we found that such checks are necessary even for data acquired by professional scientists. It must be kept in mind that thrill, motivation, and self-confidence are keys to school children engagement with science and with practical scientific activities^20,37^. Our findings that school children did no worse than untrained professional scientists in collecting ecological data (here, in estimating insect herbivory) can strengthen their confidence and help them gain motivation and a positive attitude toward science in general. Thus, even collecting and formatting the data, and sharing the process with scientists, will be valuable parts of training school children in scientific literacy. Such activities clearly contribute to our joint understanding of ecology and the nature of science in general.

## Materials and methods

### Oak selection

We designed a simple protocol that was applied by both school children and professional scientists. The protocol was written by scientists in collaboration with science instructors and communication officers. It was available in French, English, German, Spanish and Portuguese^38^.

In early 2018, each project partner selected a minimum of one mature English oak tree with lower branches accessible from the ground. Partners selected one to 18 oak trees (schoolchildren: 1-8, median = 2; scientific partners: 1-18, median = 6). We imposed no restriction on oak tree location, age or size, but professional scientists were asked to choose oaks in woods larger than 1ha. All partners measured oak tree circumference at 1.30 m from the ground and recorded oak coordinates with the GPS function of their smartphones.

All partners installed dummy caterpillars in lower branches of their own selected oak trees to estimate predation rate and haphazardly collected fresh leaves from the same trees they were expected to estimate insect herbivory. While most of the partner schools provided predation rate estimates, none undertook herbivory assessment because leaf sampling was too close to the end of the school term. We thus compared the precision and accuracy of predation rate estimates by school children *vs.* professional scientists from data collected in the ‘*Oak bodyguard*’ project framework and set up a complementary experiment to evaluate precision and accuracy of estimating insect herbivory by school children and professional scientists (see below).

### Predation rate

Six weeks after oak budburst (to control for latitudinal variation in study site locations), partners installed 20 dummy caterpillars per tree, *i.e.*, five caterpillars on each of four branches (facing north, south, east and west) and a minimum distance of 15 cm between each caterpillar. Caterpillars were made of the same green plasticine (Staedler, Noris Club 8421, green[5]) provided to all partners by the project coordinators (B. Castagneyrol, EVC). In order to standardize caterpillar size among partners, caterpillars were made from a ball of plasticine of 1 cm diameter, and gently pressed/rolled onto the middle of a 12 cm long metallic wire until a 3 cm long caterpillar was obtained. Partners were instructed to leave caterpillars on trees for 15 days prior to recording predation marks. Every caterpillar with any suspected predation marks from any guilds of potential predators (i.e., birds, mammals, reptiles and arthropods) was tagged and photographed by partner schools from three different angles to show the observed damage. Photos were labelled in such a way that file name indicated both tree and caterpillar ID. Photos were used by project leaders to double-check and standardize predation assessment made by individual partners. A second survey using the same procedure immediately followed the first one.

Although we could not organize direct training sessions with partner schools because of obvious geographical constraints, every partner received indirect training that consisted of a field ‘bite guide’. It contained a collection of photos illustrating predation marks left by different types of predators as well as ‘fake positive’ marks on plasticine surfaces by leaves, buds or nails. The different predator guilds that can be easily identified from their typical marks left on plasticine include passerine birds, rodents, snakes and lizards, molluscs and insects, mainly beetles and bush-crickets^32^. The ‘bite guide’ was available online and accessible to all partners through a hyperlink from the protocol^38^.

All partners were required to record their observations in the same standardized recording form. Partners indicated the total number of caterpillars: installed; with any type of predation marks; with no predation mark; and with predation marks left by birds (typically V-shaped beak marks and holes), arthropods (mandible marks), mammals (parallel teeth marks) or lizards (ellipse-shaped line of small teeth marks). We intentionally asked for redundant information to limit the risk of error in data reporting.

Data and biological material were collected by both school children and professional scientists during the same time period (May and July 2018). Project partners filled in the recording form and sent it to the project leader together with the photos of attacked caterpillars (partner schools) or caterpillars (professional scientists) and oak leaves. A single observer (EVC) with previous expertise in identifying predation marks on model caterpillars^39^ screened every photo to verify observations reported by partners. For each oak tree and survey period, we assessed predation rate as the proportion of dummy caterpillars with at least one predation mark. Although we asked partners to record predation marks left by different types of predators (in particular birds and arthropods), this level of precision could not be reached on photos because of low resolution. Therefore we quantified overall predation rate, regardless of predator type.

We compared predation rate as estimated in the field by school or scientific partners *vs.* predation rate as estimated afterward by a single trained observer (EVC), considered as the control treatment with paired *t*-tests. We estimated the precision and accuracy of predation rate assessments by schoolchildren and untrained professional scientists by running two separate linear mixed-effect models with predation rate estimated by school children or professional scientists as a dependent variable, predation rate estimated by a single trained professional scientist as an independent variable, and partner ID as a random factor ^33^. From each regression, we quantified the bias (that is a deviation between predation rate estimated by partners and a single trained observer) as the intercept (β_0_). Positive deviation from β_0_ = 0 indicates an overestimation of predation rate by partners. We then quantified accuracy as the regression slope (β_1_), where β_1_ = 1 indicates high accuracy and β_1_ ≠ 1 indicates that accuracy in predation rate assessment varied with actual predation rate.

### Insect herbivory

In order to compare insect herbivory estimated by school children *vs.* trained and untrained professional scientists, we set up a complementary survey (administered by AB). In April 2019, we prepared 12 sets of five oak leaves randomly drawn from a large sample of oak leaves collected in September 2018 on 162 oak trees and stored in paper bags at −18°C in a freezer. For each set of leaves, five trained professional scientists with previous experience in scoring insect herbivory on oak leaves (BC, EVC, AB, TD, YK [see acknowledgements]) estimated insect herbivory as the percentage of leaf area removed or impacted by insect herbivores by giving each individual leaf a damage score: 0: 0%, A: 1-5%, B: 6-15%, C: 16-25%, D: 26-50%, E: 51-75%, F: > 75% ^40^. In order to reduce variability in estimates of herbivory due to observers, we created digital model leaves with given amounts of simulated herbivore damage that were used as examples for the seven damage classes^38^. Leaf chewers were the main source of insect herbivory on oak leaves, but because leaves were drawn at random from a large pool of leaves, some were attacked by leaf-miners, while none had galls. We asked participants to score *total* insect herbivory, regardless of damaging agents. As a result, damage score incorporated leaf area removed by chewers as well as covered by leaf mines.

We invited 11-16 years old students (and their teachers) of six local secondary schools (equivalent US grades 6-10) to visit the first author’s research facilities (INRA research station of Pierroton, Bordeaux, France). Five groups of 10-12 students were introduced to the study of insect herbivory by the survey administrator who challenged them to score insect herbivory as accurately as professional scientists would do. Students worked in groups of 2-3. Each group was given three sets of five leaves, selected at random from the pool of 12 leaf sets. All students scored damage using the same digital model leaves as a template. In total, each of the 12 leaf sets was processed by six independent groups of students.

The same day (or the day after), we invited INRA permanent and non-permanent staff members to participate in the survey. Volunteers were researchers, engineers, technicians and MSc students. They were regarded as professional scientists, but had no previous experience in herbivory assessment (henceforth: untrained professional scientists). They received the same information from the survey administrator as secondary school students and used the same templates to score herbivory. Each of the nine volunteers processed every set of five leaves.

For each leaf set and each observer, we averaged herbivory by using the median of each damage class. Repeated handling of the same leaves may have caused some breakage, leading to a progressively increased estimation of herbivory. Due to this we first verified that herbivory did not increase with time since the first assessment. We used a linear mixed-effect model with Time (number of hours since the very first assessment) and Observer type (i.e., trained professional scientist, untrained professional scientist or student) and their interaction as a fixed effect factor and leaf set identity as a random effect factor to account for repeated measurements. We detected no effect of Time (*F*_2, 229.05_ = 0.16, *P* = 0.686) or Time × Observer interaction (*F*_2, 230.04_ = 0.43, *P* = 0.650, **Figure S1**) and therefore did not account for time in subsequent analyses.

We averaged herbivory estimates across observers belonging to the same group (i.e. trained professional scientist, untrained professional scientist or student) for each set of leaves. We first tested whether individuals with a different background differed in their estimation of insect herbivory by running linear mixed effect models with (untransformed) insect herbivory as a response variable, observer type as a fixed effect factor and leaf set identity as a random effect factor. Pairwise differences among observer types were tested by calculating contrasts among treatments. Second, we estimated the precision and accuracy of school children and untrained professional scientists as we did for predation rate (see above). Here, we note that the intercept and slope are not fully independent of each other. A steep slope within the data range may create a negative intercept, i.e. a negative value at a herbivory rate of zero, which did not occur in the data **(Figure 5b)**. Thus, the intercept and the slope should be interpreted in unison.

All analyses were done in R^41^ using packages *lmerTest* and *car*^42,43^.

## Acknowledgements

This study has been carried out with financial support from the French National Research Agency (ANR) in the frame of the Investments for the future Programme, within the Cluster of Excellence COTE (ANR-10-LABX-45). The authors warmly thank all young European citizens and their teachers who have made this study possible. They also thank professional scientists who have kindly accepted to participate in this study: Xoaquin Moreira (Misión Biológica de Galicia, MBG-CSIC), Giada Centenaro, Stefan K. Müller (Freie evangelische Schule Lörrach), Olga Mijón Pedreira (teacher IES Rosais 2, Vigo-Spain), Mikhail Kozlov and Elena Zvereva (Section of Ecology, Dept of Biology, Univ. of Turku, FI-20014 Turku, Finland), Andreas Schuldt (University of Göttingen, Forest Nature Conservation, Göttingen), Aurélien Sallé (Laboratoire de Biologie des Ligneux et des Grandes Cultures, INRA, Université d’Orléans, 45067 Orléans, France) Mickael Pihain and Andreas Prinzing (Research Unit “Ecosystèmes, Biodiversité, Evolution”, University of Rennes 1 / CNRS, 35042 Rennes, France), and Chloe Mendiondo and Claire Colliaux (Department of Agroecology, Aarhus University, Flakkebjerg Research Centre, DK-4200 Slagelse, Denmark). We thank Thomas Damestoy and Yasmine Kadiri (trained professional scientists) and Laure Dubois, Yannick Mellerin, Isabelle Lesur, Yves Ritter, Corinne Vacher, Geoffrey Haristoy, Thomas Folituu and Alex Stemmelen (untrained professional scientists) for having scored insect herbivory together with BC, EVC, AB as well as Christelle Guilloux and Ingvild Marchand (Lycée Grand Air, Arcachon), Emilie Goyran (Collège Gaston Flament, Marcheprime), Danouckka Vignal-Tudal, Erwan Thépaut (collège Leroi Gourhan, Le Bugue) and all their students.

## Contributions

BC conceived the study, analyzed the data and lead the writing. BC and EVC coordinated the research, with help from DH, MB, MKD, MMG, MSL and RT. EVC acquired and formatted the data. All authors contributed data, critically commented and edited the manuscript.

## Competing interests

The authors declare no competing financial interests.

## References

1. Kahan, D. M. et al. The polarizing impact of science literacy and numeracy on perceived climate change risks. Nat. Clim. Change 2, 732–735 (2012).

2. Fiske, S. T. & Dupree, C. Gaining trust as well as respect in communicating to motivated audiences about science topics. Proc. Natl. Acad. Sci. 111, 13593–13597 (2014).

3. Drummond, C. & Fischhoff, B. Individuals with greater science literacy and education have more polarized beliefs on controversial science topics. Proc. Natl. Acad. Sci.201704882 (2017). doi:10.1073/pnas.1704882114

4. European Commission. Green paper on citizen science. 51 (2013).

5. Haklay, M. Citizen science and policy: A European perspective. (Woodrow Wilson International Center for Scholars, 2015).

6. Wals, A. E. J., Brody, M., Dillon, J. & Stevenson, R. B. Convergence Between Science and Environmental Education. Science 344, 583–584 (2014).

7. Dickinson, J. L. et al. The current state of citizen science as a tool for ecological research and public engagement. Front. Ecol. Environ. 10, 291–297 (2012).

8. Schwartz, M. D., Betancourt, J. L. & Weltzin, J. F. From Caprio’s lilacs to the USA National Phenology Network. Front. Ecol. Environ. 10, 324–327 (2012).

9. Lucky, A. et al. Ecologists, educators, and writers collaborate with the public to assess backyard diversity in The School of Ants Project. Ecosphere 5, art78 (2014).

10. Miczajka, V. L., Klein, A.-M. & Pufal, G. Elementary School Children Contribute to Environmental Research as Citizen Scientists. PLoS ONE 10, (2015).

11. Saunders, M. E. et al. Citizen science in schools: Engaging students in research on urban habitat for pollinators. Austral Ecol. 43, 635–642 (2018).

12. Ekholm, A., Tack, A. J. M., Bolmgren, K. & Roslin, T. The forgotten season: the impact of autumn phenology on a specialist insect herbivore community on oak. Ecol. Entomol. 44, 425–435 (2019).

13. Hurlbert, A., Hayes, T., McKinnon, T. & Goforth, C. Caterpillars Count! A Citizen Science Project for Monitoring Foliage Arthropod Abundance and Phenology. Citiz. Sci. Theory Pract. 4, 1 (2019).

14. Trumbull, D., Bonney, R., Bascom, D. & Cabral, A. Thinking Scientifically during Participation in a Citizen-Science Project. Sci. Educ. 84, 265–275 (2000).

15. McLaughlin, J., Benforado, J. & Liu, S. B. Report to Congress describes the breadth and scope of Federal crowdsourcing and citizen science. citizenscience.gov (2019).

16. Makuch, K. & Aczel, M. Children and citizen science. in 391– 409 (UCL Press, 2018). doi:10.14324/111.9781787352339

17. Jenkins, L. L. Using citizen science beyond teaching science content: a strategy for making science relevant to students’ lives. Cult. Stud. Sci. Educ. 6, 501–508 (2011).

18. Koomen, M. H., Rodriguez, E., Hoffman, A., Petersen, C. & Oberhauser, K. Authentic science with citizen science and student-driven science fair projects. Sci. Educ. 102, 593–644 (2018).

19. Wells, N. M. & Lekies, K. Children and nature: following the trail to environmental attitudes and behaviour. in Citizen Science: public collaboration in environmental research 201–213 (2012).

20. Ganzevoort, W. & van den Born, R. The Thrill of Discovery: Significant Nature Experiences Among Biodiversity Citizen Scientists. Ecopsychology 11, 22–32 (2019).

21. Pandya, R. E. A framework for engaging diverse communities in citizen science in the US. Front. Ecol. Environ. 10, 314–317 (2012).

22. Jordan, R. C., Gray, S. A., Howe, D. V., Brooks, W. R. & Ehrenfeld, J. G. Knowledge Gain and Behavioral Change in Citizen-Science Programs. Conserv. Biol. 25, 1148–1154 (2011).

23. Brossard, D., Lewenstein, B. & Bonney, R. Scientific knowledge and attitude change: The impact of a citizen science project. Int. J. Sci. Educ. 27, 1099–1121 (2005).

24. Riesch, H. & Potter, C. Citizen science as seen by scientists: Methodological, epistemological and ethical dimensions. Public Underst. Sci. Bristol Engl. 23, 107–120 (2014).

25. Kelemen-Finan, J., Scheuch, M. & Winter, S. Contributions from citizen science to science education: an examination of a biodiversity citizen science project with schools in Central Europe. Int. J. Sci. Educ. 40, 2078–2098 (2018).

26. Scheuch, M. et al. Butterflies & wild bees: biology teachers’ PCK development through citizen science. J. Biol. Educ. 52, 79–88 (2018).

27. Burgess, H. et al. The science of citizen science: Exploring barriers to use as a primary research tool. Biol. Conserv. 208, (2016).

28. Ratnieks, F. L. W. et al. Data reliability in citizen science: learning curve and the effects of training method, volunteer background and experience on identification accuracy of insects visiting ivy flowers. Methods Ecol. Evol. 7, 1226–1235 (2016).

29. Steinke, D., Breton, V., Berzitis, E. & Hebert, P. D. N. The School Malaise Trap Program: Coupling educational outreach with scientific discovery. PLOS Biol. 15, e2001829 (2017).

30. Mäntylä, E. et al. From Plants to Birds: Higher Avian Predation Rates in Trees Responding to Insect Herbivory. PLoS ONE 3, e2832 (2008).

31. Roslin, T. et al. Higher predation risk for insect prey at low latitudes and elevations. Science 356, 742–744 (2017).

32. Lövei, G. L. & Ferrante, M. A review of the sentinel prey method as a way of quantifying invertebrate predation under field conditions. Insect Sci. 24, 528–542 (2017).

33. Johnson, M. T. J., Bertrand, J. A. & Turcotte, M. M. Precision and accuracy in quantifying herbivory. Ecol. Entomol. 41, 112–121 (2016).

34. Zvereva, E. L. & Kozlov, M. V. Biases in studies of spatial patterns in insect herbivory. Ecol. Monogr. in press, e01361 (2019).

35. Forstmeier, W., Wagenmakers, E.-J. & Parker, T. H. Detecting and avoiding likely false-positive findings – a practical guide. Biol. Rev. 92, 1941–1968 (2017).

36. Bayraktarov, E. et al. Do Big Unstructured Biodiversity Data Mean More Knowledge? Front. Ecol. Evol. 6, 239 (2019).

37. Ruiz-Mallen, I. et al. Citizen Science: Toward Transformative Learning. Sci. Commun. 38, 523–534 (2016).

38. Castagneyrol, B., Valdés-Correcher, E., Kaennel Dobbertin, M. & Gossner, M. Predation assessment on fake caterpillars and leaf sampling: Protocol for partner schools. protocols.io(2019). doi:10.17504/protocols.io.42pgydn

39. Valdés-Correcher, E., van Halder, I., Barbaro, L., Castagneyrol, B. & Hampe, A. Insect herbivory and avian insectivory in novel native oak forests: Divergent effects of stand size and connectivity. For. Ecol. Manag. 445, 146–153 (2019).

40. Castagneyrol, B., Giffard, B., Péré, C. & Jactel, H. Plant apparency, an overlooked driver of associational resistance to insect herbivory. J. Ecol. 101, 418–429 (2013).

41. R Core Team. R: a language and environment for statistical computing. (R fundation for statistical computing, 2018).

42. Kuznetsova, A., Brockhoff, P. B. & Christensen, R. H. B. lmerTest: Tests in Linear Mixed Effects Models. (2015).

43. Fox, J. et al. car: Companion to Applied Regression. (2016).

