## supplementary material for "Can school children support ecological research? Lessons from the *‘Oak bodyguard’* citizen science project"

- Introduction
- Materials and methods
  - Oak selection
  - Predation rate
  - Insect herbivory
- Results
  - Predation rate
  - Leaf insect herbivory
- Discussion
  - Conclusion
- Acknowledgements
- References
- Supplementary material

Bastien Castagneyrol\(^1\)\*, Elena Valdés-Correcher\(^1\), Audrey Bourdin\(^1\), Luc Barbaro\(^{2,3}\), Olivier Bouriaud\(^4\), Manuela Branco\(^5\), György Csóka\(^6\), Mihai-Leonard Duduman\(^4\), Anne-Maïmiti Dulaurent\(^7\), Csaba B. Eötvös\(^6\), Marco Ferrante\(^{8,9}\), Ágnes Fürjes-Mikó\(^6\), Andrea Galmán\(^{10}\), Martin M. Gossner\(^{11}\), Deborah Harvey\(^{12}\), Andy G. Howe\(^{13}\), Michèle Kaennel-Dobbertin\(^{11}\), Julia Koricheva\(^{12}\), Gábor L. Löveï\(^{8}\), Daniela Lupaștean\(^4\), Slobodan Milanović\(^{14,15}\), Anna Mrazova\(^{16,17}\), Lars Opgennoorth\(^{18,19}\), Juha-Matti Pitkänen\(^{20}\), Marija Popović\(^{15}\), Tomas V. Roslin\(^{20}\), Michael Scherer-Lorenzen\(^{21}\), Katerina Sam\(^{16,17}\), Marketa Tahadlova\(^{16,17}\), Rebecca Thomas\(^{12}\), Ayco J. M. Tack\(^{22}\)

**Affiliations**:

1. BIOGECO, INRA, Univ. Bordeaux, 33610 Cestas, France
2. Dynafor, INRA-INPT, University of Toulouse, Auzeville, France
3. CESCO, Museum National d’Histoire Naturelle, CNRS, Sorbonne-University, Paris, France
4. ”Ștefan cel Mare” University of Suceava, Forestry Faculty, Applied Ecology Laboratory, Universității Street 13, Suceava, Romania.
5. Centro de Estudos Florestais, Instituto Superior de Agronomia, Universidade de Lisboa;
6. NARIC Forest Research Institute, Department of Forest Protection, Hegyalja str. 18, 3232 Mátrafüred, Hungary
7. UniLaSalle, AGHYLE, UP.2018.C101, SFR Condorcet FR CNRS 3417, FR-60026 Beauvais, France
8. Department of Agroecology, Aarhus University, Flakkebjerg Research Centre, DK-4200 Slagelse, Denmark
9. Mitrani Department of Desert Ecology, Ben-Gurion University of the Negev, Midreshet Ben-Gurion, 8499000, Israel
10. Misión Biológica de Galicia (MBG-CSIC),Pontevedra, Galicia, Spain
11. Forest Entomology, Swiss Federal Research Institute WSL, Zürcherstrasse 111, CH-8903 Birmensdorf, Switzerland;
12. School of Biological Sciences, Royal Holloway University of London, Egham, UK TW20 0EX
13. Department of Geosciences and Natural Resource Management, University of Copenhagen, Denmark, Rolighedsvej 23, 1958 Frederiksberg C, Denmark
14. University of Belgrade, Faculty of Forestry, Kneza Višeslava 1, 11000 Belgrade, Serbia
15. Mendel University, Faculty of Forestry and Wood Technology, Zemedelska 3, 61 300 Brno, Czech Republic
16. Biology Centre CAS, Entomology Institute, Ceske Budejovice, 37005, Czech Republic
17. University of South Bohemia, Faculty of Science, Ceske Budejovice, 37005, Czech Republic;
18. Swiss Federal Research Institute WSL, Biodiversity and Conservation Biology, Ecological Genetics, Zürcherstrasse 111, 8903 Birmensdorf
19. Department of Ecology, Philipps-Universität Marburg, Karl-von-Frisch Strasse 8, 35043 Marburg
20. Department of Ecology, Spatial Foodweb Ecology Group, Department of Agricultural Sciences, PO Box 27 (Latokartanonkaari 5), FI-00014 University of Helsinki, Finland
21. Geobotany, Faculty of Biology, University of Freiburg, Schaenzlestr. 1, 79104 Freiburg, Germany
22. Department of Ecology, Environment and Plant Sciences, Stockholm University, SE-106 91 Stockholm, Sweden

**Contributions**: BC conceived the study, analyzed the data and lead the writing. BC and EVC coordinated the research, with help from DH, MB, MKD, MMG, MSL and RT. EVC acquired and formatted the data. All authors contributed data, critically commented and edited the manuscript.

**Competing interests** :The authors declare no competing financial interests.

---

**Abstract -** Scientific knowledge in the field of ecology is increasingly enriched bydata acquired by the general public participating in citizen science (CS) programs. Yet, doubts remain about the reliability of such data, in particular when acquired by schoolchildren. We built upon an ongoing CS program - *Oak bodyguards* - to assess the ability of European schoolchildren to accurately estimate the strength of biotic interactions in terrestrial ecosystems. We used standardized protocols to estimate predation rates on artificial caterpillars and insect herbivory on oak leaves and compared estimates made by schoolchildren, trained and untrained professional scientists (with no or limited expertise in predation or herbivory assessment). Compared to trained scientists, both schoolchildren and untrained professional scientists overestimated predation rates, but assessments made by the latter were more consistent. schoolchildren overestimated insect herbivory, as did untrained professional scientists. Thus, raw data acquired by schoolchildren participating in CS programs cannot be used and require several quality checks. However, such data are of no less value than data collected by untrained professional scientists and can be calibrated for bias.

**Keywords -** Artificial larvae, Citizen science, Data quality, Insect herbivory, Measurement bias

---

### Introduction

Scientific knowledge is more accessible than ever before, particularly due to an increase in open-access publications and outreach activities of scientists worldwide. Still, many science topics in life and environmental sciences are highly controversial in society, even among individuals with substantial science literacy and education (Kahan et al. 2012, Fiske and Dupree 2014, Drummond and Fischhoff 2017). Citizen science (CS) programs rely on participation of the general public in scientific research in collaboration with or under the direction of professional scientists (European Commission 2013, Haklay 2015). The rapid development of these programs, in addition to vastly increasing available data, offers an unprecedented opportunity to bridge gaps between science and society, by engaging the general public with the process of science, and increasing motivation and interest in scientific topics.

Citizen science programs in the field of ecology can benefit both science and society (Wals et al. 2014). For professional scientists, involving the general public enables the collection of data on broader spatial and temporal scales than would otherwise be possible (i.e., ‘crowdsourcing’). This practice has been recognized as a highly effective way to track various biological phenomena (Dickinson et al. 2012, Schwartz et al. 2012). Typical CS studies in ecology address the effect of environmental factors on biodiversity (e.g. Lucky et al. 2014, Miczajka et al. 2015, Saunders et al. 2018) or climate change impact on plant or animal phenology (Schwartz et al. 2012, Ekholm et al. 2019, Hurlbert et al. 2019). In turn, volunteers engaged in CS programs can gain recognition for their skills and develop a deeper understanding of scientific concepts and the scientific process (Trumbull et al. 2000). This may positively contribute to both science and environmental education (Wals et al. 2014) and raise awareness of environmental issues. As a result, CS programs are now promoted by major funding agencies in Europe and North America (e.g., European Commission 2013, McLaughlin et al. 2019).

Engaging schoolchildren and their teachers can enhance the long-term educational and social goals of CS programs, for several reasons (Makuch and Aczel 2018). First, school pupils are guided by their instructors when learning about the scientific question raised by the CS program, as well as about the nature and social aspects of science (Jenkins 2011, Koomen et al. 2018). Second, exposure to outdoor nature during childhood increases motivation and provides a long-lasting positive relationship with the environment while increasing people’s knowledge about nature (Wells and Lekies 2012, Ganzevoort and van den Born 2019). Third, targeting schoolchildren for CS projects has the potential to engage a wider cross-section of society in science (Wells et al. 2015) than other CS projects involving self-selecting volunteers (they choose whether to be involved which can lead to the underrepresentation of many social groups, although strategies exist to increase engagement, Pandya 2012).

Nonetheless, the enthusiastic views of win-win interactions through CS programs have been questioned by social scientists and ecologists (Jordan et al. 2011). The former point out that the educational and social impact may be overstated (Trumbull et al. 2000, Brossard et al. 2005, Riesch and Potter 2014, Kelemen-Finan et al. 2018, Scheuch et al. 2018), while the latter are concerned about the accuracy of data collected by the general public (Burgess et al. 2016), especially when schoolchildren are involved. The main reason for these concerns is that CS data are arguably of lower quality than those collected by professional scientists (Riesch and Potter 2014, Burgess et al. 2016, Makuch and Aczel 2018). It has been proposed that data collected by schoolchildren involved in CS programs can contribute to environmental research, provided that research methods are kept simple and require skills that the children already have or are able to gain when mentored by adults (Miczajka et al. 2015, Makuch and Aczel 2018, Saunders et al. 2018), and that the participant receives training, even remotely (Ratnieks et al. 2016). However, only a few studies have directly compared the quality of data acquired by professional scientists vs. the general public (Miczajka et al. 2015, Steinke et al. 2017, Saunders et al. 2018). Quantitative evaluations of the impact of CS programs on science are thus needed to reciprocally engage citizens with science, and scientists with citizens.

For each leaf set and each observer, we averaged herbivory by using the median of each damage class. Repeated handling of the same leaves may have caused some breakage, leading to a progressively increased estimation of herbivory. Due to this we first verified that herbivory did not increase with time since the first assessment. We used a linear mixed-effect model with *Time* (number of hours since the very first assessment) and *Observer type* (i.e., trained professional scientist, untrained professional scientist or student) and their interaction as a fixed effect factor and leaf set identity as a random effect factor to account for repeated measurements. We detected no effect of *Time* (*df* = (1, 229.05), *F* = 0.16, *P* = 0.686) or *Time* \(\times\) *Observer* interaction (*df* = (2, 227.15), *F* = 0.431, *P* = 0.65, **Fig. S1**) and therefore did not account for time in subsequent analyses.

We averaged herbivory estimates across observers belonging to the same group (i.e. trained professional scientist, untrained professional scientist or student) for each set of leaves. We first tested whether individuals with a different background differed in their estimation of insect herbivory by running linear mixed effect models with (untransformed) insect herbivory as a response variable, observer type as a fixed effect factor and leaf set identity as a random effect factor. Pairwise differences among observer types were tested by calculating contrasts among treatments. Second, we estimated the precision and accuracy of schoolchildren and untrained professional scientists as we did for predation rate (see above). Here, we note that the intercept and slope are not fully independent of each other. A steep slope within the data range may create a negative intercept, *i.e.* a negative value at a herbivory rate of zero, which did not occur in the data **(Fig. 5b)**. Thus, the intercept and the slope should be interpreted in unison.

All analyses were done in *R* (R Core Team 2018) using packages `lmerTest` and `car` (Kuznetsova et al. 2015, Fox et al. 2016).

### Results

#### Predation rate

In total, 5520 dummy caterpillars were installed on 153 oak trees by 34 partner schools in 8 countries and 28 scientific partners in 14 countries throughout Europe **(Figure 1)**. Dummy caterpillars were exposed on trees for 16 days on average (range: 3 - 56). Among the 1775 dummy caterpillars installed by schoolchildren, 640 were identified by EVC with attack marks by predators (*i.e.*, 36.06 %). Among the 3745 dummy caterpillars installed by professional scientists, 1268 were found to be attacked by predators (*i.e.*, 33.86 %).

> Figure 1. Location of oak trees included in the study.

Both schoolchildren (paired *t*test: *t* = -6.31, *P* = 0) and professional scientists (*t* = -3.8, *P* = 0) overestimated predation rate as compared to estimates made by a single trained observer **(Figure 2)**. Detailed examination of pairwise comparisons at the tree level confirmed that partner schools consistently overestimated predation rates as compared to a single observer, while the sign of deviation in scoring by scientific partners was more balanced between over- and under-estimation **(Figure 2)**.

> Figure 2. Comparison of predation rate estimated by schoolchildren or professional scientists vs. a single trained observer (EVC: Elena Valdés-Correcher). Points joined by grey lines represent the same tree.

Predation rate estimates by schoolchildren was more biased (intercept estimate \(\pm\) SE: \(\beta\_0\) = 41.16 \(\pm\) 8.38) than those by professional scientists (\(\beta\_0\) = 23.42 \(\pm\) 5.17). Likewise, schoolchildren made less accurate predation assessments (slope estimate: \(\beta\_1\) = 0.42 \(\pm\) 0.23) than professional scientists did (\(\beta\_1\) = 0.66 \(\pm\) 0.06, **Figure 4**).

> Figure 3. Precision and accuracy of schoolchildren (A) and professional scientists (B) in assessing predation rate (% artificial larvae with predation marks). Dots represent predation rate aggregated at the level of oak trees for each survey separately. Dashed lines indicate a 1:1 relation. Bold blue lines represent linear regression slopes of predation rate by schoolchildren or professional scientists on assessments made by a single trained observer (EVC: Elena Valdés-Correcher). Regression equations: \(y =\) 0.42 \(\times\) \(x\) + 41.16, marginal (fixed effects) \(R\_m^2\) = 0.07, conditional (fixed plus random effects) \(R\_c^2\) = 0.53 (A) and \(y =\) 0.66 \(\times\) \(x\) + 23.42, \(R\_m^2\) = 0.31, \(R\_c^2\) = 0.79 (B).

#### Leaf insect herbivory

Insect herbivory estimates by trained professional scientists were the lowest (mean \(\pm\) SE: 9 \(\pm\) 0.51 %, range: 2.2 - 19.6%, **Figure S2**), whereas insect herbivory estimates by untrained professional scientists were the highest (14.65 \(\pm\) 1.01 %, range: 3.8 - 62%, **Figure S1**). schoolchildren estimates of insect herbivory were intermediate (11.55 \(\pm\) 0.64 %, range 2.2 - 27.4%, **Figure S2**). Thus, both untrained professional scientists and schoolchildren consistently overestimated insect herbivory compared to trained professional scientists (\(F\_{2,22}\) = 27.31, *P* < 0.001, **Figs 4 and S2**). Interestingly, untrained professional scientists overestimated insect herbivory compared to both schoolchildren and trained professional scientists **(Figure 4)**.

> Figure 4. Pairwise comparisons between insect herbivory as estimated by schoolchildren, trained and untrained professional scientists. Boxplots represent mean insect herbivory aggregated across observers at the level of individual leaf sets. Thick grey lines connect dots corresponding to the same leaf sets. Different letters above boxplots indicate significant differences between treatments.

Herbivory estimated by schoolchildren was less biased (intercept estimate \(\pm\) SE: \(\beta\_0\) = 0.84 \(\pm\) 1.58) than that of untrained professional scientists (\(\beta\_0\) = -3.5 \(\pm\) 2.61). Likewise, schoolchildren made more accurate herbivory assessments (slope estimate: \(\beta\_1\) = 1.19 \(\pm\) 0.17) than untrained professional scientists did (\(\beta\_1\) = 1.99 \(\pm\) 0.27, **Figure 5**). Note that on average, unlike schoolchildren, untrained professional scientists overestimated herbivory across the observed range of herbivory. The negative intercept for untrained professional scientists here is driven by a slope estimate close to \(\beta\_1 = 2\).

**How can we make data collected by schoolchildren more reliable?** Citizen science programs can help to generate a large amount of data, but the quality has been questioned, especially when these ‘big data’ are not well structured by standard protocols (Burgess et al. 2016, Bayraktarov et al. 2019). Few studies have evaluated the quality of data collected by schoolchildren participating in citizen science programs (Miczajka et al. 2015, Steinke et al. 2017, Saunders et al. 2018). It emerges from these studies that schoolchildren can actually provide data accurate enough to support ecological research, provided that the tasks they are requested to undertake are adapted to their skills and that they receive proper training (Miczajka et al. 2015, Ratnieks et al. 2016, Saunders et al. 2018). Although we could not provide face-to-face training sessions for every school partner involved in the ‘*Oak bodyguards*’ project, the project methodology was simple and based on a detailed protocol. Nonetheless, this simplicity did not suffice to guarantee unbiased data, as illustrated by the fact that schoolchildren consistently overestimated predation rates. We therefore emphasize that citizen science programs relying on data collected by schoolchildren should include several checks of data quality and appropriate mitigation procedures. In particular, training sessions undertaken face-to-face or at least remotely must be planned before data collection (Ratnieks et al. 2016). Finally, whenever possible, the researcher analyzing the data should recover the raw material collected by children, or at the very least pictures allowing the re-assessment of measurements (Steinke et al. 2017, Ekholm et al. 2019). Importantly, these recommendations also hold true for large multi-partners research programs, as we also detected bias in data collected by professional scientists (Zvereva and Kozlov 2019). Whether variability in observations made by schoolchildren is random or can be modelled using appropriate covariates is an important question deserving further attention.

#### Conclusion

Here we found that schoolchildren involved in CS programs can support ecological research, but only if their contributions are considered with caution. The acquisition of reliable data requires experimental procedures that are easy to implement, but even so, a measurement of interpretation bias seems essential. Several quality checks and curation procedures are therefore needed prior to using data collected by schoolchildren for ecological research. Unexpectedly, we found that such checks are necessary even for data acquired by professional scientists. It must be kept in mind that thrill, motivation, and self-confidence are keys to schoolchildren engagement with science and with practical scientific activities (Ruiz-Mallen et al. 2016, Ganzevoort and van den Born 2019). Our findings that schoolchildren did no worse than untrained professional scientists in collecting ecological data (here, in estimating insect herbivory) can strengthen their confidence and help them gain motivation and a positive attitude toward science in general. Thus, even collecting and formatting the data, formatting them, and sharing the process with scientists, will be valuable parts of training schoolchildren in scientific literacy. Such activities clearly contribute to our joint understanding of ecology and the nature of science in general.

### Acknowledgements

This study has been carried out with financial support from the French National Research Agency (ANR) in the frame of the Investments for the future Programme, within the Cluster of Excellence COTE (ANR-10-LABX-45). The authors warmly thank all young European citizens and their teachers who have made this study possible. They also thank professional scientists who have kindly accepted to participate in this study: Xoaquin Moreira (Misión Biológica de Galicia, MBG-CSIC), Giada Centenaro, Stefan K. Müller (Freie evangelische Schule Lörrach), Olga Mijón Pedreira (teacher IES Rosais 2, Vigo-Spain), Mikhail Kozlov and Elena Zvereva (Section of Ecology, Dept of Biology, Univ. of Turku, FI-20014 Turku, Finland), Andreas Schuldt (University of Göttingen, Forest Nature Conservation, Göttingen), Aurélien Sallé (Laboratoire de Biologie des Ligneux et des Grandes Cultures, INRA, Université d’Orléans, 45067 Orléans, France) Mickael Pihain and Andreas Prinzing (Research Unit “Ecosystèmes, Biodiversité, Evolution”, University of Rennes 1 / CNRS, 35042 Rennes, France), and Chloe Mendiondo and Claire Colliaux (Department of Agroecology, Aarhus University, Flakkebjerg Research Centre, DK-4200 Slagelse, Denmark). We thank Thomas Damestoy (TD) and Yasmine Kadiri (YK) (trained professional scientists) and Laure Dubois, Yannick Mellerin, Isabelle Lesur, Yves Ritter, Corinne Vacher, Geoffrey Haristoy, Thomas Folituu and Alex Stemmelen (untrained professional scientists) for having scored insect herbivory together with BC, EVC, AB as well as Christelle Guilloux and Ingvild Marchand (Lycée Grand Air, Arcachon), Emilie Goyran (Collège Gaston Flament, Marcheprime), Danouckka Vignal-Tudal, Erwan Thépaut (collège Leroi Gourhan, Le Bugue) and all their students.

### References

- Bayraktarov, E., G. Ehmke, J. O’Connor, E. L. Burns, H. A. Nguyen, L. McRae, H. P. Possingham, and D. B. Lindenmayer. 2019. Do Big Unstructured Biodiversity Data Mean More Knowledge? Frontiers in Ecology and Evolution 6:239.
- Brossard, D., B. Lewenstein, and R. Bonney. 2005. Scientific knowledge and attitude change: The impact of a citizen science project. International Journal of Science Education 27:1099–1121.
- Burgess, H., L. DeBey, H. Froehlich, N. Schmidt, J. Hille Ris Lambers, J. Tewksbury, and J. K. Parrish. 2016. The science of citizen science: Exploring barriers to use as a primary research tool. Biological Conservation 208.
- Castagneyrol, B., B. Giffard, C. Péré, and H. Jactel. 2013. Plant apparency, an overlooked driver of associational resistance to insect herbivory. Journal of Ecology 101:418–429.
- Castagneyrol, B., E. Valdés-Correcher, M. Kaennel Dobbertin, and M. Gossner. 2019. Predation assessment on fake caterpillars and leaf sampling: Protocol for partner schools. protocols.io.
- Dickinson, J. L., J. Shirk, D. Bonter, R. Bonney, R. L. Crain, J. Martin, T. Phillips, and K. Purcell. 2012. The current state of citizen science as a tool for ecological research and public engagement. Frontiers in Ecology and the Environment 10:291–297.
- Drummond, C., and B. Fischhoff. 2017. Individuals with greater science literacy and education have more polarized beliefs on controversial science topics. Proceedings of the National Academy of Sciences:201704882.
- Ekholm, A., A. J. M. Tack, K. Bolmgren, and T. Roslin. 2019. The forgotten season: the impact of autumn phenology on a specialist insect herbivore community on oak. Ecological Entomology 44:425–435.
- European Commission. 2013. Green paper on citizen science. Page 51.
- Fiske, S. T., and C. Dupree. 2014. Gaining trust as well as respect in communicating to motivated audiences about science topics. Proceedings of the National Academy of Sciences 111:13593–13597.
- Forstmeier, W., E.-J. Wagenmakers, and T. H. Parker. 2017. Detecting and avoiding likely false-positive findings – a practical guide. Biological Reviews 92:1941–1968.
- Fox, J., S. Weisberg, D. Adler, D. Bates, G. Baud-Bovy, S. Ellison, D. Firth, M. Friendly, G. Gorjanc, S. Graves, R. Heiberger, R. Laboissiere, G. Monette, D. Murdoch, H. Nilsson, D. Ogle, B. Ripley, W. Venables, D. Winsemius, A. Zeileis, and R-Core. 2016. car: Companion to Applied Regression.
- Ganzevoort, W., and R. van den Born. 2019. The Thrill of Discovery: Significant Nature Experiences Among Biodiversity Citizen Scientists. Ecopsychology 11:22–32.
- Haklay, M. 2015. Citizen science and policy: A European perspective. Woodrow Wilson International Center for Scholars, Washington, DC.
- Hurlbert, A., T. Hayes, T. McKinnon, and C. Goforth. 2019. Caterpillars Count! A Citizen Science Project for Monitoring Foliage Arthropod Abundance and Phenology. Citizen Science: Theory and Practice 4:1.
- Jenkins, L. L. 2011. Using citizen science beyond teaching science content: a strategy for making science relevant to students’ lives. Cultural Studies of Science Education 6:501–508.
- Johnson, M. T. J., J. A. Bertrand, and M. M. Turcotte. 2016. Precision and accuracy in quantifying herbivory. Ecological Entomology 41:112–121.
- Jordan, R. C., S. A. Gray, D. V. Howe, W. R. Brooks, and J. G. Ehrenfeld. 2011. Knowledge Gain and Behavioral Change in Citizen-Science Programs. Conservation Biology 25:1148–1154.
- Kahan, D. M., E. Peters, M. Wittlin, P. Slovic, L. L. Ouellette, D. Braman, and G. Mandel. 2012. The polarizing impact of science literacy and numeracy on perceived climate change risks. Nature Climate Change 2:732–735.
- Kelemen-Finan, J., M. Scheuch, and S. Winter. 2018. Contributions from citizen science to science education: an examination of a biodiversity citizen science project with schools in Central Europe. International Journal of Science Education 40:2078–2098.
- Koomen, M. H., E. Rodriguez, A. Hoffman, C. Petersen, and K. Oberhauser. 2018. Authentic science with citizen science and student-driven science fair projects. Science Education 102:593–644.
- Kuznetsova, A., P. B. Brockhoff, and R. H. B. Christensen. 2015. lmerTest: Tests in Linear Mixed Effects Models.
- Lövei, G. L., and M. Ferrante. 2017. A review of the sentinel prey method as a way of quantifying invertebrate predation under field conditions. Insect Science 24:528–542.
- Lucky, A., A. M. Savage, L. M. Nichols, C. Castracani, L. Shell, D. A. Grasso, A. Mori, and R. R. Dunn. 2014. Ecologists, educators, and writers collaborate with the public to assess backyard diversity in The School of Ants Project. Ecosphere 5:art78.
- Makuch, K., and M. Aczel. 2018. Children and citizen science. Pages 391–409. UCL Press.
- Mäntylä, E., G. A. Alessio, J. D. Blande, J. Heijari, J. K. Holopainen, T. Laaksonen, P. Piirtola, and T. Klemola. 2008. From Plants to Birds: Higher Avian Predation Rates in Trees Responding to Insect Herbivory. PLoS ONE 3:e2832.
- McLaughlin, J., J. Benforado, and S. B. Liu. 2019, June 18. Report to Congress describes the breadth and scope of Federal crowdsourcing and citizen science.
- Miczajka, V. L., A.-M. Klein, and G. Pufal. 2015. Elementary schoolchildren Contribute to Environmental Research as Citizen Scientists. PLoS ONE 10.
- Pandya, R. E. 2012. A framework for engaging diverse communities in citizen science in the US. Frontiers in Ecology and the Environment 10:314–317.
- R Core Team. 2018. R: a language and environment for statistical computing. R fundation for statistical computing, Vienna, Austria.
- Ratnieks, F. L. W., F. Schrell, R. C. Sheppard, E. Brown, O. E. Bristow, and M. Garbuzov. 2016. Data reliability in citizen science: learning curve and the effects of training method, volunteer background and experience on identification accuracy of insects visiting ivy flowers. Methods in Ecology and Evolution 7:1226–1235.
- Riesch, H., and C. Potter. 2014. Citizen science as seen by scientists: Methodological, epistemological and ethical dimensions. Public Understanding of Science (Bristol, England) 23:107–120.
- Roslin, T., B. Hardwick, V. Novotny, W. K. Petry, N. R. Andrew, A. Asmus, I. C. Barrio, Y. Basset, A. L. Boesing, T. C. Bonebrake, E. K. Cameron, W. Dáttilo, D. A. Donoso, P. Drozd, C. L. Gray, D. S. Hik, S. J. Hill, T. Hopkins, S. Huang, B. Koane, B. Laird-Hopkins, L. Laukkanen, O. T. Lewis, S. Milne, I. Mwesige, A. Nakamura, C. S. Nell, E. Nichols, A. Prokurat, K. Sam, N. M. Schmidt, A. Slade, V. Slade, A. Suchanková, T. Teder, S. van Nouhuys, V. Vandvik, A. Weissflog, V. Zhukovich, and E. M. Slade. 2017. Higher predation risk for insect prey at low latitudes and elevations. Science 356:742–744.
- Ruiz-Mallen, I., L. Riboli-Sasco, C. Ribrault, M. Heras, D. Laguna, and L. Perie. 2016. Citizen Science: Toward Transformative Learning. Science Communication 38:523–534.
- Saunders, M. E., E. Roger, W. L. Geary, F. Meredith, D. J. Welbourne, A. Bako, E. Canavan, F. Herro, C. Herron, O. Hung, M. Kunstler, J. Lin, N. Ludlow, M. Paton, S. Salt, T. Simpson, A. Wang, N. Zimmerman, K. B. Drews, H. F. Dawson, L. W. J. Martin, J. B. Sutton, C. C. Webber, A. L. Ritchie, L. D. Berns, B. A. Winch, H. R. Reeves, E. C. McLennan, J. M. Gardner, C. G. Butler, E. I. Sutton, M. M. Couttie, J. B. Hildebrand, I. A. Blackney, J. A. Forsyth, D. M. Keating, and A. T. Moles. 2018. Citizen science in schools: Engaging students in research on urban habitat for pollinators. Austral Ecology 43:635–642.
- Scheuch, M., T. Panhuber, S. Winter, J. Kelemen-Finan, M. Bardy-Durchhalter, and S. Kapelari. 2018. Butterflies & wild bees: biology teachers’ PCK development through citizen science. Journal of Biological Education 52:79–88.
- Schwartz, M. D., J. L. Betancourt, and J. F. Weltzin. 2012. From Caprio’s lilacs to the USA National Phenology Network. Frontiers in Ecology and the Environment 10:324–327.
- Steinke, D., V. Breton, E. Berzitis, and P. D. N. Hebert. 2017. The School Malaise Trap Program: Coupling educational outreach with scientific discovery. PLOS Biology 15:e2001829.
- Trumbull, D., R. Bonney, D. Bascom, and A. Cabral. 2000. Thinking Scientifically during Participation in a Citizen-Science Project. Science Education 84:265–275.
- Valdés-Correcher, E., I. van Halder, L. Barbaro, B. Castagneyrol, and A. Hampe. 2019. Insect herbivory and avian insectivory in novel native oak forests: Divergent effects of stand size and connectivity. Forest Ecology and Management 445:146–153.
- Wals, A. E. J., M. Brody, J. Dillon, and R. B. Stevenson. 2014. Convergence Between Science and Environmental Education. Science 344:583–584.
- Wells, N. M., and K. Lekies. 2012. Children and nature: following the trail to environmental attitudes and behaviour. Pages 201–213 Citizen Science: public collaboration in environmental research. Cornell University Press. Ithaca, NY.
- Wells, N. M., B. M. Myers, L. E. Todd, K. Barale, B. Gaolach, G. Ferenz, M. Aitken, C. R. Henderson, C. Tse, K. O. Pattison, C. Taylor, L. Connerly, J. B. Carson, A. Z. Gensemer, N. K. Franz, and E. Falk. 2015. The Effects of School Gardens on Children’s Science Knowledge: A Randomized Controlled Trial of Low-Income Elementary Schools. International Journal of Science Education 37:2858–2878.
- Zvereva, E. L., and M. V. Kozlov. 2019. Biases in studies of spatial patterns in insect herbivory. Ecological Monographs in press:e01361.

### Supplementary material

> Figure S1. Effect of the duration of the experiment on leaf insect herbivory. The x-axis represents times (number of hours) since herbivory was assessed for the first time. Dots represent herbivory as estimated by each observer on individual leaf sets.

> Figure S2. Comparison of leaf insect herbivory assessments by schoolchildren, trained and untrained professional scientists for each set of five leaves. Leaf samples (A-L) have been ordered according to leaf insect herbivory as estimated by trained professional scientists. Dots and error bars represent means and SE, for data aggregated across observers.

Bastien Castagneyrol

1 mai 2019
